# Calculating Protein-Ligand Residence Times Through State Predictive Information Bottleneck based Enhanced Sampling

**DOI:** 10.1101/2024.04.16.589710

**Authors:** Suemin Lee, Dedi Wang, Markus A. Seeliger, Pratyush Tiwary

**Affiliations:** Biophysics Program and Institute for Physical Science and Technology, University of Maryland, College Park 20742, USA; Department of Pharmacological Sciences, Stony Brook University, Stony Brook, NY 11794-8651, USA; Department of Chemistry and Biochemistry and Institute for Physical Science and Technology, University of Maryland, College Park 20742, USA; University of Maryland Institute for Health Computing, Rockville, United States

## Abstract

Understanding drug residence times in target proteins is key to improving drug efficacy and understanding target recognition in biochemistry. While drug residence time is just as important as binding affinity, atomic-level understanding of drug residence times through molecular dynamics (MD) simulations has been difficult primarily due to the extremely long timescales. Recent advances in rare event sampling have allowed us to reach these timescales, yet predicting protein-ligand residence times remains a significant challenge. Here we present a semi-automated protocol to calculate the ligand residence times across 12 orders of magnitudes of timescales. In our proposed framework, we integrate a deep learning-based method, the state predictive information bottleneck (SPIB), to learn an approximate reaction coordinate (RC) and use it to guide the enhanced sampling method metadynamics. We demonstrate the performance of our algorithm by applying it to six different protein-ligand complexes with available benchmark residence times, including the dissociation of the widely studied anti-cancer drug Imatinib (Gleevec) from both wild-type Abl kinase and drug-resistant mutants. We show how our protocol can recover quantitatively accurate residence times, potentially opening avenues for deeper insights into drug development possibilities and ligand recognition mechanisms.

**TOC Graphic:** 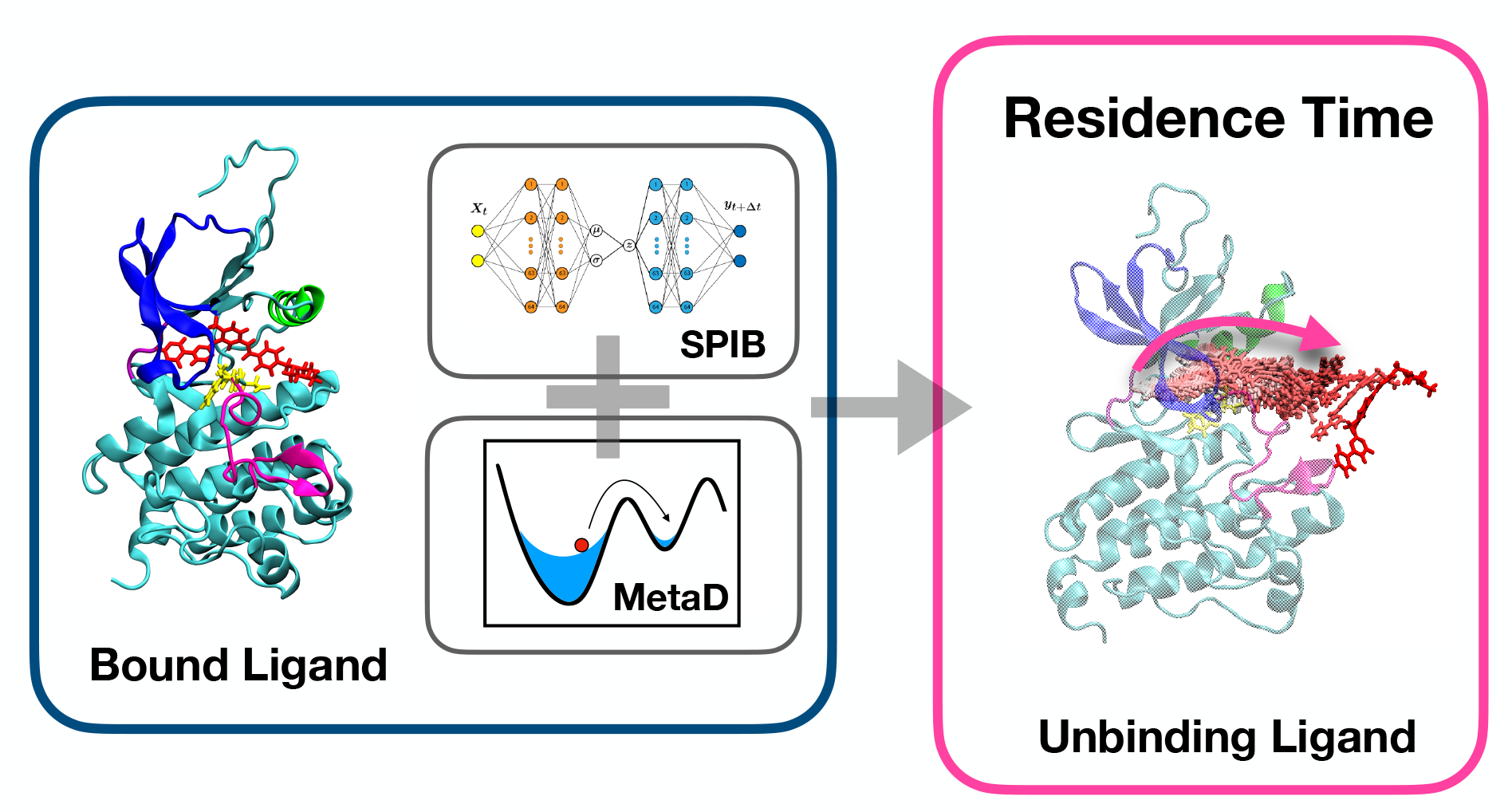

## I. INTRODUCTION

In drug discovery and fundamental biochemistry, accurately understanding the thermodynamics and the kinetics of the drug dissociation process from the target protein is an important task for improving drug efficacy and analyzing biomolecular target recognition.^1,2^ Traditionally, the main emphasis has been given to the binding affinity, which is indeed an important indicator for determining drug efficiency.^3,4^ However, through numerous studies, it has been demonstrated that for drug efficacy and safety, considering the residence time, which is the duration a drug remains bound to its target protein, and is often defined by the reciprocal of the dissociation rate (*k*_off_), is as if not more important than binding affinity alone.^5–11^ For instance, studies of Abl kinase domain resistance to the drug Imatinib showed that mutations with the same binding affinity resulted in a greater reduction in Imatinib efficacy due to changes in residence time, suggesting that drug residence time plays an important role in drug efficacy.^9^ Furthermore, other studies have shown that extended residence time allows pharmacodynamic effects to persist even after the concentration in circulation drops.^11^ The relevance of the residence time is restricted not just to drug efficacy but more generally to the chemistry of different processes in human health and disease.^12^

However, exploring the time-dependent interactions underlying protein-ligand complexes has been a challenge, both in experimental and computational studies. Experimentally probing these encounters limitations in capturing dynamical characteristics such as transition states. Furthermore, it is difficult to perform experiments that can directly capture the drug dissociation pathways and interactions in atomic resolution. Using molecular dynamics (MD) simulations could provide such an atomic-level insight, but examining the kinetics through MD has been challenging due to extremely long timescales. Namely, while drug dissociation occurs at a timescales of hours or slower, MD is limited to around microseconds to milliseconds even with state-ofthe-art specialized supercomputers.^13^ Recent progress in the development of rare event sampling has allowed circumventing this timescale limitation by making it possible to observe ligand dissociation with full atomistic resolution.^14–21^ Through accelerated sampling of the conformation space, these methods can, more or less in an semi-automated manner effectively, reach the timescales that were unattainable in MD simulations.^10^ While these rare event methods come in a wide range of flavors and implementations, a generic overarching theme for methods that can measure rare event kinetics is (i) the choice of a few biasing or progress coordinates.^22,23^ (ii) a protocol to enhance the sampling by using these coordinates,(iii) calculate the unbiased true kinetics from the simulations, and while not possible in all rare event methods, (iv)calculate the reliability of the kinetics previously calculated. Despite the success in this area of using rare event methods to accelerate sampling in MD, predicting the kinetics of protein-ligand dissociation with advanced MD simulations in an semi-automated and robust manner continues to be significantly challenging.^24^ The challenges include but are not limited to each of the four steps mentioned in the previous paragraph.

To address this issue, in this manuscript we propose a semi-automated protocol to study protein-ligand residence times. This method streamlines all four steps above, enabling the recovery of quantitatively accurate determination of residence times for various proteinligand complexes across 12 order of magnitude in residence times. In our proposed framework, we integrate a deep learning-based method for learning an approximate reaction coordinate (RC) with metadynamics, a widely used enhanced sampling approach. The approximate one- or two-dimensional RC captures the slow degrees of freedom relevant to the ligand dissociation process. It then serves as the biasing variable for enhanced sampling, and also enables visualization of the underlying dissociation mechanisms and transition pathways. For the first key part of our protocol, which is to learn the RC, here we use the State Predictive Information Bottleneck (SPIB) approach, which is a member of the Reweighted Autoencoded Variational Bayes for Enhanced Sampling (RAVE) class of methods.^25,26^ For a generic protein-ligand complex, dissociation is generally not a 2-state process.^27^ In addition to the bound and unbound states, one can expect an *a priori* unknown number of metastable states lying on the dominant dissociation pathway. One of the key advantages of SPIB is that it enables learning the number of metastable states and their approximate locations. SPIB expresses the RC as a shallow or deep neural network as a function of selected input features, which are protein ligand contact maps in this work.

The second key part of our protocol involves metadynamics based enhanced sampling. Like any deep learning framework, SPIB needs data for its training. We initiate our approach with trial runs using independent well-tempered metadynamics^16^ that specifically target hydrogen bond breakage of key interactions in the crystal structure. We perform many such runs in parallel, biasing different hydrogen bonds, and select only those for which dissociation is observed for further analysis. We perform SPIB on these input trajectories to learn an approximate RC, which is biased in a new round of well-tempered metadynamics. Continuing from the RAVE framework, this protocol continues until no further enhancements in the effectiveness of RC are detected. Finally, we perform multiple independent infrequent metadynamics (iMetaD) simulations biasing along the SPIB RC to calculate residence time.^28^ The reliability of the kinetics so obtained from infrequent metadynamics is calculated using the Kolmogorov–Smirnov test from Ref.29

We demonstrate our algorithm’s performance on different systems, comparing against much longer unbiased MD simulations when possible and also *in vitro* experiments previously reported. Together, these systems span 12 orders of magnitude in timescales, ranging from nanoseconds to minutes. We start with simple systems, such as the dissociation of two fragments with millimolar binding affinity from the protein FKBP, which we can compare with long unbiased MD.^30,31^ This is followed by the study of benzene dissociation from T4 Lysozyme, which we can compare against the previously obtained experimental results.^32^ Finally, we study the Abl kinase-Imatinib system, where Imatinib (commercially sold as Gleevec) is recognized for its significant efficacy in treating cancer leukemia.^33,34^ Understanding the dissociation process in this system is crucial for gaining insights into how the drug interacts with its target protein.^10^ For Imatinib as well, our semi-automated protocol is able to provide an accurate estimation of residence times against Abl kinase in its wild type (WT) and two drug resistant N368S and L364I mutant forms.^9^

The structure of this paper is outlined as follows: In Section II, we introduce our systematic protocol for computing ligand dissociation or residence time, achieved by integrating deep neural networks with enhanced sampling. Section III provides the outcomes of diverse applications concerning the protein-ligand residence times. Lastly, Section IV summarizes our findings and discusses future directions. An open source code demonstrating the full protocol has been provided at GitHub.

## II. METHODS

In this section, we provide the background behind all methods used in this work. This includes metadynamics in the two flavors of well-tempered metadynamics and infrequent metadynamics, and the SPIB approach. We further provide an overview of the protocol where we integrate metadynamics with SPIB in a seamless fashion to recover reliable kinetics of protein-ligand dissociation. Finally, we provide details of the MD setup for the different systems studied here.

### A. Well-tempered and infrequent metadynamics

In metadynamics, a history-dependent bias potential is constructed as the simulation progresses. This bias is built as a Gaussian function that depends on carefully chosen collective variables, which should encapsulate relevant slow degrees of freedom. This helps the system escape metastable states and explore new parts of the free energy landscape. In the well-tempered variant^14^, the height of the Gaussian is gradually decreased as a function of simulation time. The product of this procedure is the free energy surface as a function of the originally biased collective variables or any generic variables, even if not directly biased through a reweighting procedure.^35^

In infrequent metadynamics,^28^ the frequency at which bias is added is reduced so that the possibility of adding bias as the simulation is moving across the transition state (TS) becomes low. Naturally, with more infrequent bias deposition, one would approach the limit of unbiased MD, but this would also mean that there is no acceleration of configurational sampling. Under suitable approximations, which we describe shortly, the unbiased kinetics can be recovered by scaling the biased timescale with the following acceleration factor,^36,37^

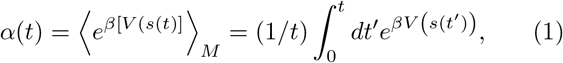

where *V* (*s, t*^*′*^) is the metadynamics time-dependent bias, *β* = 1*/k*_*B*_*T* the inverse temperature multiplied by the Boltzmann constant *k*_*B*_, and subscript *M* indicates averaging over metadynamics run time of *t*. We refer to the adjusted timescale, 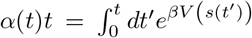, as the accelerated time. The reliability of the timescales so obtained can be quantitatively assessed through a Kolmogorov–Smirnov (KS) test.^29^ A p-value greater than 0.05 in the Kolmogorov-Smirnov (KS) test indicates that the residence time does not significantly deviate from the Poisson process model expected for rare events such as protein-ligand dissociation. This suggests that the accelerated timescales are reliable.

There are two underlying assumptions for infrequent metadynamics to be reliable and useful: (1) the system displays timescale separation, comprising extended amounts of time spent in metastable states, and relatively little time during the actual transition events crossing between states, and (2) one can construct low-dimensional variables that capture the metastable dynamics experienced by the system in its full dimensionality. The first assumption depends on the system itself and will be met as long as there is fundamental underlying separation of timescales, expected as long as the system is not too disordered.^38^ In the next sub-section, we further elaborate on the second assumption, and demonstrate how it is met using SPIB.

### B. State Predictive Information Bottleneck (SPIB)

To recover an accurate estimation of kinetics using infrequent metadynamics, one needs an approximate low-dimensional RC that preserves the metastable dynamics experienced by the system in its full higher dimensions. To address this concern in a data-driven and system-agnostic manner, different deep-learning approaches have been proposed recently.^22,39^ One of these is the Reweighted Autoencoded Variational Bayes for Enhanced Sampling (RAVE)^26^ framework. Here, we use RAVE in an improved formulation for learning the RC expressed as a state predictive information bottleneck (SPIB).^25^ SPIB has proven successful in learning the RC for diverse applications from crystal nucleation to ligand permeation through lipids and protein conformational change.^40–43^

The key idea in SPIB is to learn the RC as a lowdimensional variable that is most predictive about the system’s future metastable state while using the minimal information from the system’s current configuration. More technically, SPIB training involves processing raw trajectory input data to predict future states of the system after a time delay Δ*t*. This input information is compressed into a bottleneck variable *χ* by passing through an encoder neural network that minimizes mutual information shared between the bottleneck and the input. At the same time, the information from the bottleneck is passed through a decoder-type network that predicts the future state of the system, maximizing the mutual information between the bottleneck and the future state. The number of metastable states, as well as their locations in the higher-dimensional input space, are learned on-the-fly by assuming an appropriate mixture of Gaussians prior for the bottleneck variable *χ*. Through this tradeoff between minimizing and maximizing two mutual information, the process identifies metastable states and learns the approximate RC as the bottleneck variable *χ*. For the problem of protein-ligand dissociation, if Δ*t* is significantly large, then SPIB will consider the entire configuration space as a single metastable state. In the context of protein-ligand systems, it will regard both the bound and unbound conditions as a single state. As Δ*t* is reduced, SPIB learns more states, such as the bound state and unbound state as well as any metastable states lying on the dissociation pathway.

## C. Overall pipeline

In this work, we demonstrate our technique for six different proteins-ligand complexes. These are FKBP-DSS and FKBP-DMSO, T4 lysozyme L99A mutant-Benzene, and Abl kinase WT-Imatinib with two more mutants of Abl kinase N368S and L364I.^10,31^ These systems have experimentally validated ligand residence times ranging from ∼ 10 ns to ∼ 10^3^ s. The details of each system preparation are described in the SI. We begin with preliminary runs of well-tempered metadynamics by biasing along the trial variables such as hydrogen bonds or, more generally, the distances between the ligand and the protein. When the hydrogen bonds existed in the system, there were approximately 1 to 4 were accessible in our given applications. In cases where a hydrogen bond was absent between the protein and ligand, such as in T4 Lysozyme L99A-benzene, we selected the distances between adjacent binding pocket helices and the center of mass (COM) of the ligand as an alternative. In this step, we carefully selected trajectories where the ligand had successfully dissociated from the protein, making them suitable for SPIB training trajectory data. Initially, we simulated 4 trajectories for FKBP-ligand and 8 and 10 trajectories for T4 lysozyme and kinase-Imatinib, respectively, for two trajectories per each trial variable. The FKBP ligand fully dissociated within 50 ns of simulations. For T4 Lysozyme, 4 out of 10 trajectories stayed bound within 100 ns. For kinase-Imatinib, 4 out of 8 trajectories remained bound within 200 ns. Consequently, these trapped trajectories were excluded from SPIB training data set.

We now describe the order parameters (OPs) that we used as input features for training SPIB. These OPs comprised the pair-wise distance between the COM of the ligand and every third residue of the C*α* atom of the protein, without including flexible terminal regions. Here, the number of input OPs needs to be taken with caution. We find that choosing too many OPs can increase the complexity of the RC learned by SPIB, thus destabilizing the metadynamics biasing process. Heuristically, we find that the protocol in this work is most stable for less than 100 OPs. While we acknowledge that a more advanced feature selection could further improve this process, we deliberately chose simple and straightforward distance metrics to be our input features as a proof-of-principle and to demonstrate the simplicity and efficacy of the framework. Upon appropriate selection of the OPs, we proceed with SPIB training.

We learn the RC as a two-dimensional linear encoder expressed as a function of the input OPs. Unlike other previous studies^10,31^, here we consistently use a twodimensional RC instead of one-dimensional. While the 2-d RC is slower during subsequent metadynamics, it is more reliable in encapsulating the metastability relevant to drug dissociation. For each round, we perform 8 independent well-tempered metadynamics runs using the learned RC. Runs resulting in dissociation are combined for the subsequent round of SPIB training. While these are biased runs, we consciously ignore the bias while training the SPIB. For high-energy metastable states, considering the bias through reweighting can often make those states appear as noise due to their low Boltzmann weights. We iterate between such rounds of well-tempered metadynamics and training the SPIB, until the acceleration factor from Eq.1 no longer decreases. We emphasize that the accelerated timescales from well-tempered metadynamics are not indicative of true kinetics but serve to quantify the effectiveness of the RC in facilitating dissociation with minimal bias. Once we no longer observed improvement in the accelerated time from rounds of well-tempered metadynamics, we finalize the RC and switch to infrequent metadynamics. More precisely, we perform roughly 10-12 independent trials of infrequent metadynamics from the most optimized RC that has the shortest average accelerated time.

This entire workflow is illustrated in Fig. 1. This protocol was applied to all the protein-ligand systems in this work. If any modifications were made to this workflow at any steps or to the system, the details are discussed in the following Sec.III.

**FIG. 1:**
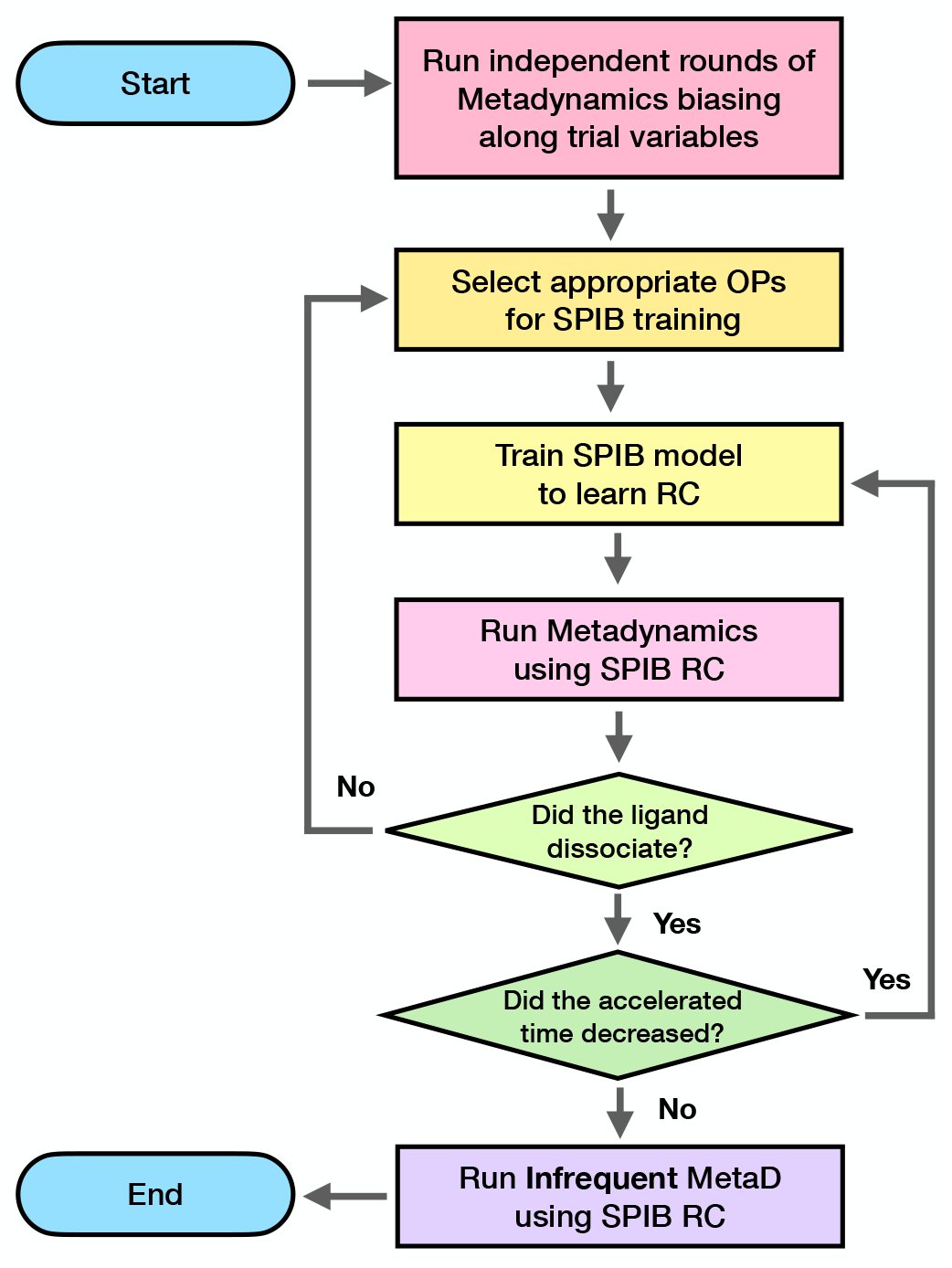
Flow chart illustrating the automated SPIB-metadynamics protocol in this work for obtaining residence times of protein-ligand complexes. First, we start with initial trial runs of well-tempered metadynamics biasing along trial OPs as described in Sec.II. We iteratively apply SPIB and metadynamics multiple times until the SPIB RC is quantified through accelerated time reduction. Finally, we run infrequent metadynamics biasing along the converged RC for an accurate estimation of the residence time.

**FIG. 2:**
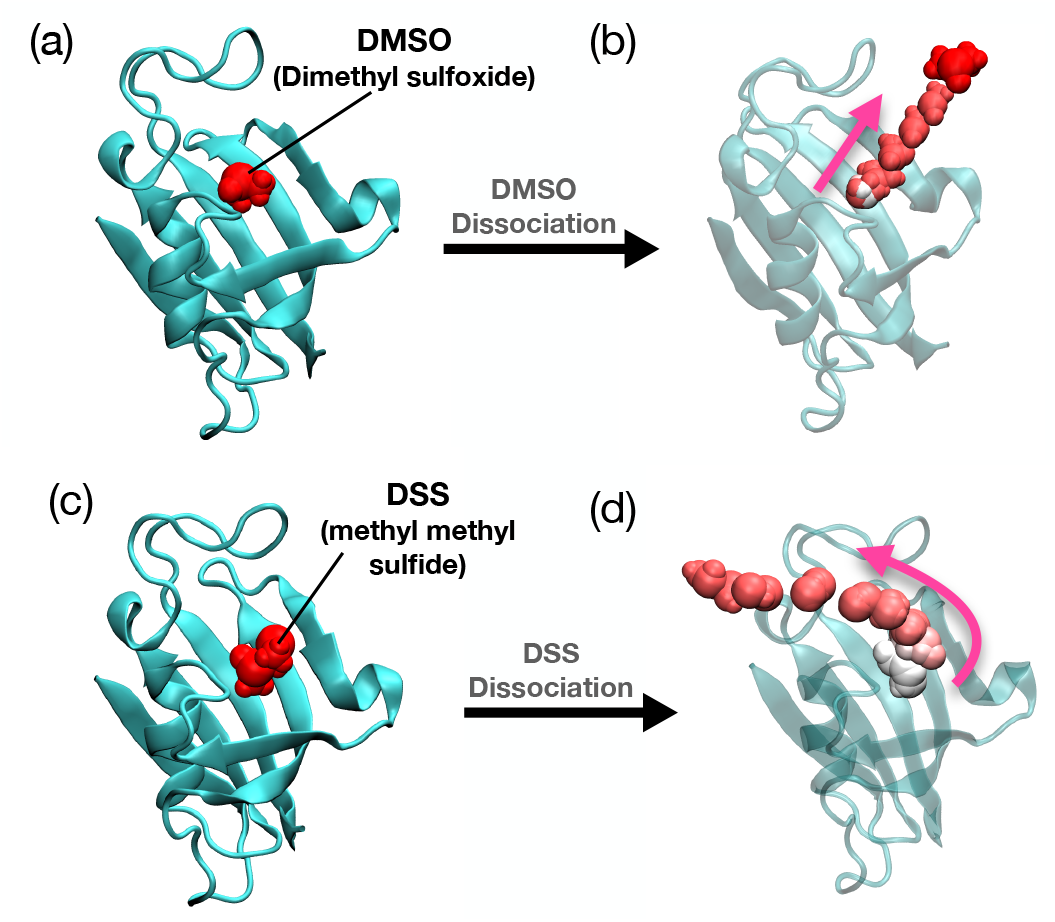
(a) FKBP with DMSO bound system (PDB ID:1D7H). The ligand DMSO is depicted in red. (b) The dissociation pathway for DMSO is illustrated in a color gradient, starting with white for the initial time steps, transitioning to pink for the intermediate stage, and red to signify the final time steps. (c) FKBP with DSS bound system (PDB ID:1D7I). The ligand DSS is depicted in red. (d) The dissociation pathway for DSS is illustrated in a color gradient, starting with white for the initial time steps, transitioning to pink for the intermediate stage, and red to signify the final time steps.

## D. Simulation setup

The simulations were performed using GROMACS 2019.4^44^ patched with PLUMED version 2.6.^45^ The protein was prepared by CHARMM36 force field^46^ and all the ligands were prepared with CHARMM General Force Field (CGenFF).^47^ The simulations were performed under constant number, pressure, and temperature (NPT) conditions that were set up at 303.15 K temperature and 1 bar pressure with the Nosé-Hoover thermostat^48,49^ and Parrinello-Rahman barostat^50^ respectively. Long-range electrostatics were calculated using the particle-mesh Ewald (PME) method.^51^ The MD integration timestep was 2 fs. The systems were solvated with TIP3P water models and equilibrated under the NPT ensemble for 5 ns on Abl Kinase system and T4 lysozyme system. The FKBP-ligand complex was equilibrated for only 100 ps to prevent ligand dissociation during the equilibration step. Further details of the system preparation are discussed in the SI.

## III. RESULTS

### A. FKBP with DMSO and DSS

FK506 binding protein (FKBP) belongs to a large family of proteins frequently observed in eukaryotes and is closely linked to various human diseases. In particular, due to the millimolar binding affinity of the ligand to the FKBP and the small size of the protein, it is a well-studied system through different computational methods.^20,30,52,53^ In this study, we applied our protocol to FKBP using two different ligands, dimethyl sulfoxide (DMSO) and methyl (methylsulfinyl) methyl sulfide (DSS), calculating the residence time for each ligand. These FKBP-ligand complexes are known to exhibit a relatively short residence time in the tens of nanosecond regime, which makes it possible to benchmark residence times with unbiased simulation results.

We applied our protocol, as outlined in Sec. II and illustrated in the flowchart in Fig. 1, without additional modifications. In the initial metadynamics runs, we biased the system along the single hydrogen bond formed between residue Ile56 in FKBP and the ligand —the sole hydrogen bond in the system. Following the first stage, we selected 26 order parameters (OPs) around the ligand’s binding pocket and conducted 2 rounds and 1 round of SPIB stage to achieve minimal accelerated time for DMSO and DSS ligands, respectively. Further details of the reduction in accelerated time according to the SPIB stages can be found in the SI Fig.S1 (a) and (b).

The cumulative results show a residence time of 29 ns with a p-value of 0.36 for the FKBP-DMSO system and 210 ns with a p-value of 0.07 for the FKBP-DSS system, meeting the 0.05 threshold for the reliability of dynamics from metadynamics proposed in Ref. 29 and subsequently also used in protein-ligand dissociation studies.^17^ While preserving the relative trends for DMSO and DSS, this represents a two to three-fold difference compared to our unbiased simulation benchmarks. The KS test results and the curve fit plots can be found in the SI Fig. S3 (a) and (b).

### B. T4 Lysozyme L99A with benzene

We now focus on T4 Lysozyme, an enzyme from the T4 bacteriophage, crucial in virus replication by breaking down cell walls for virus particle release.^54^ In particular, we examined the L99A mutant, known for its easier ligand release compared to the WT.^32,55^ Our chosen ligand was benzene. We replicated the standard protocol from Sec.II and Fig. 1, except for one modification on the selection of trial variables in the trial run. This was needed because the ligand benzene does not possess any hydrogen bond donors or acceptors.

We choose our trial variables to be the distances between the centers of mass (COM) of five helix residue domains (helices C, D, E, F, and G) and the COM of the ligand benzene. The five helices shown in Fig. 3(b) were chosen for the initial simulation run because they are the closest domains that are in contact with the bound benzene, and we can assume these helices will contribute the most to the dissociation. After the first stage of our initial trial, we again selected our OPs as the pair-wise distance between the ligand and the protein residues in the binding pocket of benzene. Subsequently, we conduct 3 rounds of SPIB well-tempered metadynamics to achieve minimal accelerated time and the result can be found in SI Fig. 1 (c). This setup allowed us to investigate the benzene release pathway and its residence time. In Ref. 56, the benzene dissociation process was studied with the weighted ensemble approach^57^ and paths were explicitly differentiated depending on their dissociation pathways. While we also observed various dissociation paths, we did not apply path clustering and considered them all collectively. Although we did not implement path clustering methods, it can be done as suggested for instance in Ref. 58, using a dynamic time-warping algorithm. We collectively combined all the independent 12 rounds of infrequent metadynamics and found that the benzene residence time was 1.3 ms with a p-value of 0.38. The KS test results and the curve fit plots can be found in the SI Fig. S3 (c). Experimentally, the benzene dissociation rate from T4 lysozyme L99A using NMR spectroscopy has been reported as 1.3*±*0.3 s (*k*_off_ = 800 *±* 200 s^−1^ at 303K).^32^ This indicates that our calculated value is within the experimental range.

**FIG. 3:**
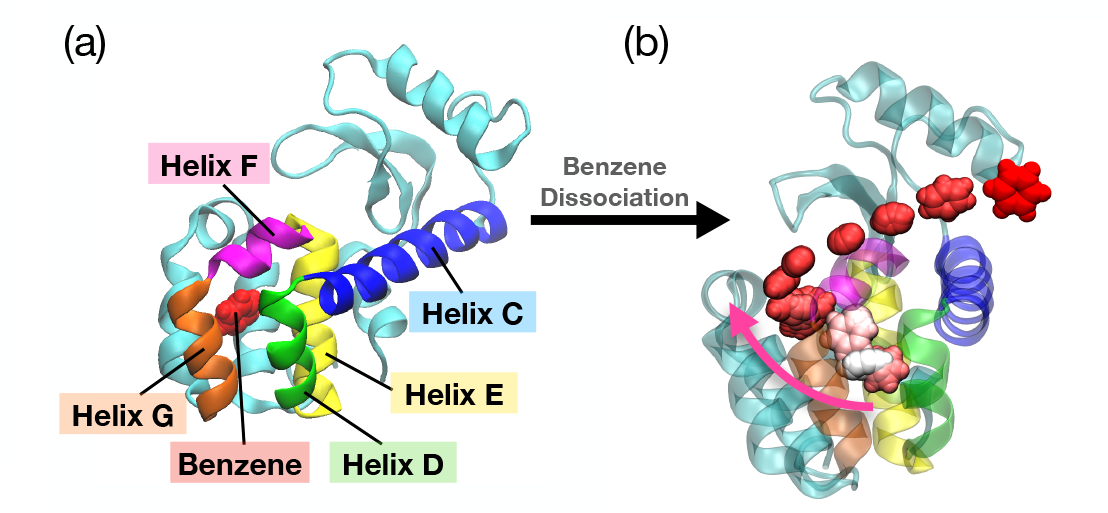
T4 Lysozyme L99A mutant with benzene bound system. (a) The structure (PDB ID: 1L83) of the T4 Lysozyme L99A, with the helix domains that are used in the initial trials of simulations depicted in different colors. Helix C is colored in blue, helix D is colored in green, helix E is colored in yellow, helix F is colored in magenta, and helix G is colored in orange. The ligand benzene, shown in red, is bound in the binding pocket of the T4 Lysozyme. (b) The dissociation pathway for benzene is illustrated in a color gradient, starting with white for the initial time steps, transitioning to pink for the intermediate stage, and red to signify the final time steps.

**FIG. 4:**
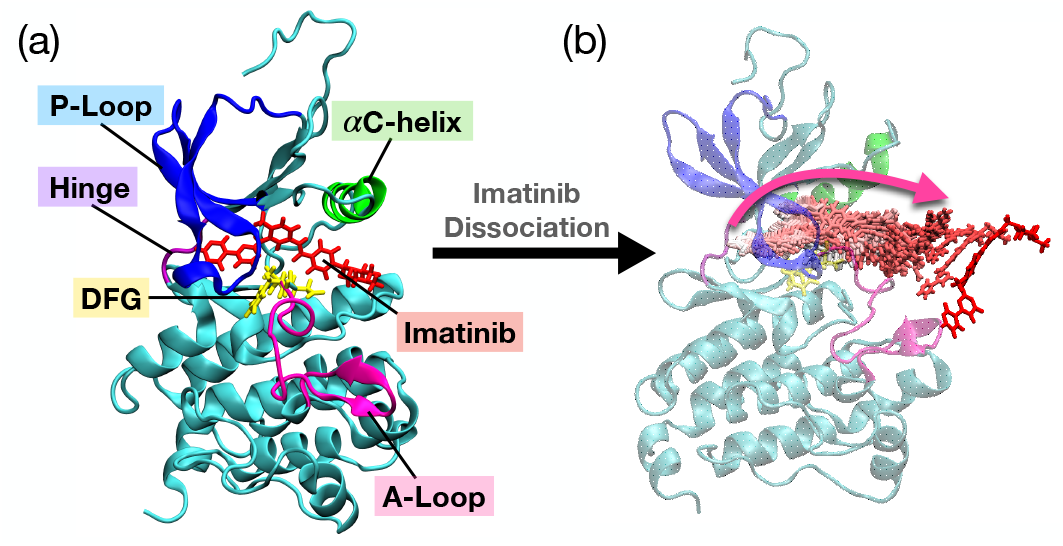
Wild Type (WT) Abl-kinase with Imatinib bound system (PDB ID: 1OPJ). (a) Important structural features are labeled in different colors. P-loop is colored in blue, *α*C-Helix is colored in green, hinge is colored in purple, DFG motif is colored in yellow and A-loop is colored in pink. Imatinib is shown in red and bound to the ATP binding pocket of kinase. (b) One of the dissociation pathways for Imatinib is illustrated in a color gradient. Starting with white for the initial time steps, transitioning to pink for the intermediate stage, and red to signify the final time steps.

### C. Abl Kinase Domain with Imatinib

Abl tyrosine kinase is an enzyme that plays a key role in cytokine signaling, and its excessive activity can lead to leukemia. For the final test of our protocol in this work, we move to an even more challenging system dissociation of the anti-cancer drug Imatinib from Abl kinase in its WT and two mutations, N368S and L364I. We again replicated our protocol to WT and N368S mutant kinase, as outlined in Sec.II and Fig. 1 without additional modifications. We applied the same protocol to L364I mutant kinase, but one modification was made to the OP selection, which is explained later. In the initial trial runs of metadynamics, we biased the system along all 4 hydrogen bonds that are formed between the Imatinib and kinase protein involving residues Glu286, Thr315, Met318 and Asp381. After the first stage, we again chose our OPs as the pair-wise distance between every third residue of the kinase and the COM of the Imatinib. This OP selection was applied to the WT kinase and N368S mutant kinase, which allowed the ligand to dissociate successfully during the metadynamics runs after the SPIB training stage. However, the same choice of OPs for the L364I mutant failed to make Imatinib dissociate from the kinase within 200 ns of simulation time. Thus, instead of the pair-wise distance, we chose the same 4 hydrogen bond features that were used at our first stage of initial trial runs of metadynamics as our OPs for training SPIB. Such a back-up protocol is easy to implement in future applications of SPIB-iMetaD. We conducted 3 rounds, 2 rounds, and 4 rounds of the SPIB stage for WT, N368S mutants, and L364I mutants, respectively, for reduction of accelerated time. The detailed results of the reduction in accelerated time according to the SPIB stages can be found in the SI Fig.S2.

Once we achieve minimal accelerated time, we switch to infrequent metadynamics and perform 20-30 independent simulation rounds per kinase system, which is higher than the number of runs performed for previous systems. This is because we observed various dissociation pathways, and as previously found in Ref. 10, each pathway is associated with different dissociation times, possibly explaining kinetics-driven resistance to Imatinib in patients treated with it. Using our clustering mechanism of the dissociation pathways, we only report the residence time of pathways with p-values exceeding 0.05 here. Results with p-values smaller than 0.05, deemed less convincing, were not presented but can further be found in the SI Table S1.

Through systematic analysis, we found that the residence time for WT was 1150 s with a p-value of 0.07, and for mutant N368S, it was 330 s with a p-value of 0.19. Both kinases have been underestimated compared to the experimental values. However, within the uncertainty error, both results show good agreement with the experimental results. The L364I mutant, on the other hand, had a residence time of 10000 s with a p-value of 0.06, which was overestimated by 14-fold. This may have been caused by the different selection of OPs for the SPIB training stage, or from not properly accounting for pathway heterogeneity, or from other sources of error, including those in experiments and in the forcefield. We acknowledge that advanced feature selection and path clustering approaches could have further improved our results. The KS test results and the curve fit plots can be found in the SI Fig. S5.

## D. Viewing all systems together

Our results for all systems are summarized in Table I and in Fig. 5. In Figure 5, the results show a good overall agreement between the benchmark and the simulation within the error bars across twelve orders of magnitude. Zooming in, we can see that the differences are within an order of magnitude. The FKBP system results showed a threefold difference compared to our unbiased simulation benchmark, while T4 lysozyme closely matched the experimental results. For Abl kinase, we also observed good agreement with the experimental results for WT and N368S, but L364I showed a sixfold difference compared to the experiments due to possible reasons as speculated in Sec. III C. While there were some discrepancies, the majority of the systems demonstrated a good alignment within the uncertainty error. This alignment suggests that our semi-automated protocol, which integrates SPIB with metadynamics, was able to identify the optimized RC and improve the accuracy of residence time calculations.

**TABLE 1:**
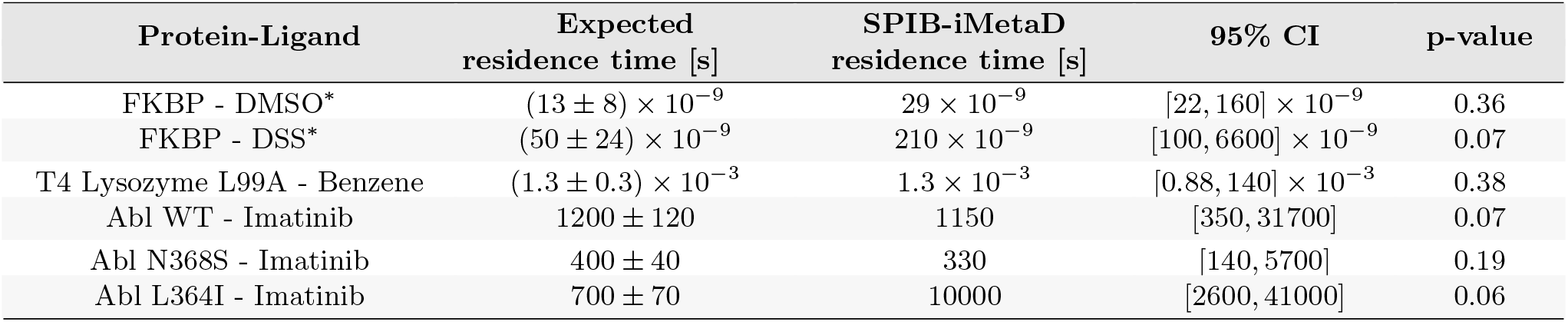
Summary table for all experimental/unbiased simulation results (expected residence time) and simulation results (SPIB-iMetaD residence time) from our own automated protocol approach, including p-values. The asterisk (*) indicates unbiased simulation results, while the experimental results are shown without asterisks. 95% CI was reported for the SPIB-iMetaD residence time measurements. The expected residence time for FKBP-DMSO and FKBP-DSS results are from our own unbiased simulation. The expected residence time for T4 Lyzozyme L99A-Benzene is from in Ref. 32 measured using NMR spectroscopy. The expected residence times for Abl Kinase of WT, N368S, and L364I-Imatinib are from Ref. 9 measured using NanoBRET.

| Protein-Ligand | Expected residence time [s] | SPIB-iMetaD residence time [s] | 95% CI | p-value |
| --- | --- | --- | --- | --- |
| FKBP - DMSO* | $(13 \pm 8) \times 10^{-9}$ | $29 \times 10^{-9}$ | $[22, 160] \times 10^{-9}$ | 0.36 |
| FKBP - DSS* | $(50 \pm 24) \times 10^{-9}$ | $210 \times 10^{-9}$ | $[100, 6600] \times 10^{-9}$ | 0.07 |
| T4 Lysozyme L99A - Benzene | $(1.3 \pm 0.3) \times 10^{-3}$ | $1.3 \times 10^{-3}$ | $[0.88, 140] \times 10^{-3}$ | 0.38 |
| Abl WT - Imatinib | $1200 \pm 120$ | 1150 | [350, 31700] | 0.07 |
| Abl N368S - Imatinib | $400 \pm 40$ | 330 | [140, 5700] | 0.19 |
| Abl L364I - Imatinib | $700 \pm 70$ | 10000 | [2600, 41000] | 0.06 |

**FIG. 5:**
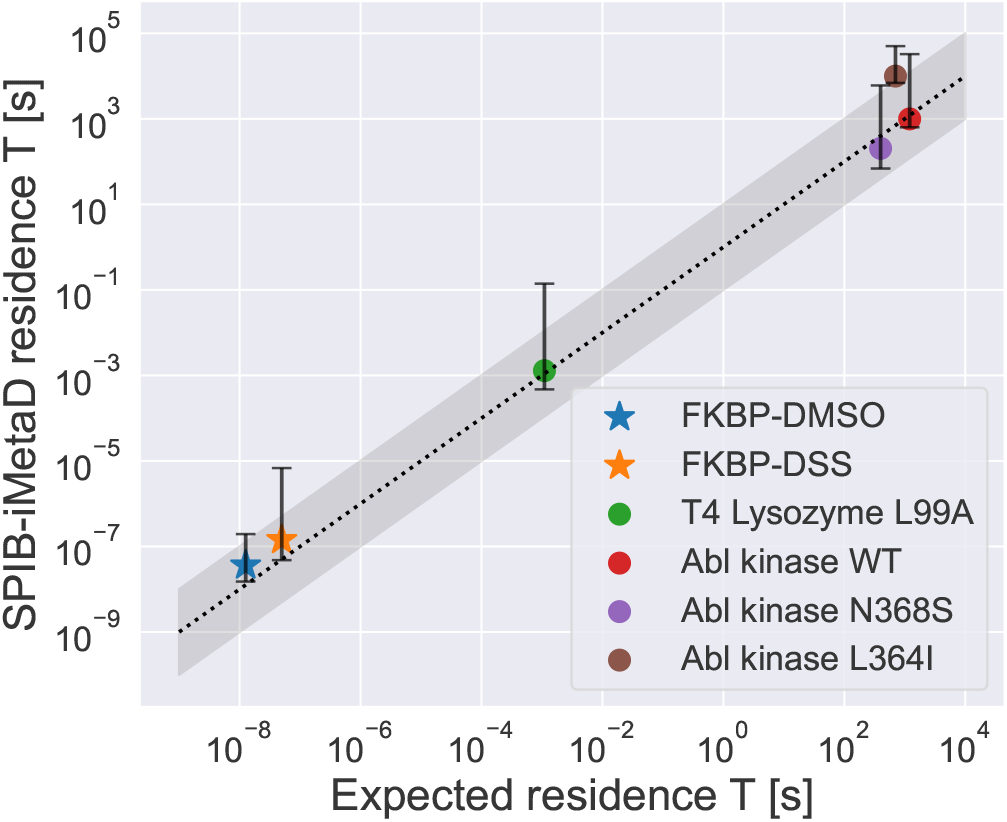
Summary graph of all the experimental/unbiased simulation results (expected residence time) versus simulation results (SPIB-iMetaD residence time) from our own semi-automated protocol approach. The circle symbol signifies the utilization of experimental data for the expected residence time, while the star symbol denotes the use of unbiased simulation results. The error bar represents 95% confidence interval computed using the bootstrapping method. The grey-shaded region indicates the 1-order of magnitude boundary along the linear dotted line.

## IV. CONCLUSION

In this study, we have introduced a semi-automated protocol to calculate residence times for a variety of protein-ligand complexes over a wide range of timescales. Our proposed framework integrates a deep learningbased method SPIB to learn reaction coordinate (RC) with the enhanced sampling approach of metadynamics. The protocol begins with the initial trial runs of well-tempered metadynamics biasing along trial variables, such as hydrogen bonds, for each protein-ligand system. Once the ligand has successfully dissociated from the protein, the pair-wise distance between the ligand COM and protein from the given trajectory is used as our input features for training SPIB. We iterate between such rounds of metadynamics and training SPIB to refine RC until the acceleration factor no longer decreases. Subsequently, we run independent rounds of infrequent metadynamics on the SPIB learned RC, where the acceleration factor has been minimized, to accurately determine the residence time. Following this, we estimated the dissociation rate through curve-fitting and evaluated the reliability of the results by the KS-test. Particularly, we demonstrated our protocol for six different proteins-ligand complexes that are FKBP-DSS and FKBP-DMSO, T4 lysozyme L99A-benzene, and Abl kinase-Imatinib, including WT and its two mutants. These systems successfully showed good agreement between our simulation results and experimental/unbiased simulation benchmark.

Despite the consistency between benchmarks on residence times and the SPIB-iMetaD simulations, challenges remain that need to be addressed and can be further improved in the future. First of all, it would be worthwhile to implement a more automated approach to cluster different pathways and obtain path-specific residence times following for instance the protocol from Ref. 58. Second, although we intentionally chose simple distance metrics to our OPs as input features for SPIB, we empirically learned that despite the same choice of OPs for different Abl kinase mutants, it could lead to the non-dissociation of ligands. In this case, as a simple alternative, we chose the hydrogen bond distances as OPs. However, we acknowledge that the appropriate selection of OPs that serve as input features to SPIB can be further improved through advanced feature selection. Third, reducing computational time is important. One limitation of our work, especially in large-scale protein-ligand systems, lies in the computational expense requirements. Effectively reducing this time can improve efficacy and make this application more feasible in a variety of systems, including protein-ligand systems and protein-protein complexes. Finally, while this work combined RC learned from SPIB with infrequent metadynamics to obtain the residence times, this could be replaced with any of the new variants of infrequent metadynamics^59,60^ Our ultimate goal is to adapt this protocol to a variety of protein-ligand complexes and other kinetics methods, enhance our understanding of the dissociation pathway, and learn important features of drug dissociation from reliable RC space.

## Supporting information

Detailed simulation setups for FKBP-ligand, T4 Lysozyme L99A-benzene, and Abl kinase-Imatinib systems are available in the supporting information (SI). This SI also includes rounds of accelerated time reduction across SPIB-metadynamics iterations, results of each Kolmogorov-Smirnov (KS) test with corresponding curve fit plots, and the path clustering approach for the Abl-kinase domain.

## Acknowledgements

This work was supported by NIH/NIGMS under award number R35GM142719 (P.T.) and R35GM119437 (M.A.S.). We thank UMD HPC’s Zaratan and NSF

ACCESS (project CHE180027P) for computational resources. P.T. is an investigator at the University of Maryland-Institute for Health Computing, which is supported by funding from Montgomery County, Maryland and The University of Maryland Strategic Partnership: MPowering the State, a formal collaboration between the University of Maryland, College Park and the University of Maryland, Baltimore. We would like to thank Mirnal Shekhar for assisting in the preparation of the Abl kinase simulation files, Zack Smith and Akashnathan Aranganathan for insightful discussions, and Eric Beyerle and Ruiyu Wang for proofreading our manuscript.

## Code availability statement

A Python code for semi-automated protocol and demo on the FKBP-DMSO system is available on GitHub (www.github.com/tiwarylab/SPIB_kinetics). For the demo version, we set up the simulations using OpenMM^61^, instead of GROMACS^44^. The results of this paper were performed using GROMACS^44^, as discussed in Sec. II D. This online version is for demonstration purposes only illustrating the key steps underlying our protocol.

## Conflict of Interest

The authors declare the following competing financial interest(s): P.T. is a consultant to Schrödinger, Inc. and is on their Scientific Advisory Board.

## Notes

### Competing Interest Statement

The authors declare the following competing financial interest(s): P.T. is a consultant to Schrodinger, Inc. and is on their Scientific Advisory Board.

