## Supplementary Materials for "Calculating Protein-Ligand Residence Times Through State Predictive Information Bottleneck based Enhanced Sampling"

### 1 Systems Preparations

#### FKBP-ligand

Molecular dynamics (MD) simulations were initiated using the crystal structures of DMSO bound to FKBP protein (PDB ID:1D7H) and DSS bound to FKBP protein (PDB ID:1D7I). The setup for these simulations was prepared by the CHARMM-GUI [1] solution builder module. We included the sodium ($Na^+$) and chloride ($Cl^-$) ions at an ionic concentration of 100 mM, along with the addition of TIP3P water molecules [2] for solvation. This procedure was uniformly applied to both the FKBP-DMSO- and FKBP-DSS complexes, ensuring consistency in simulation preparation. For the FKBP-DMSO system, a total of 16 $Na^+$ ions, 17 $Cl^-$ ions and $\sim$ 9k TIP3 water molecules were included in the system. Similarly, for the FKBP-DSS system, a total of 15 $Na^+$ ions, 16 $Cl^-$ ions and $\sim$ 8k TIP3 water molecules were included in the system. By solvating the structures in a TIP3P water box, one edge distance was extended to $\sim 65\mathring{A}$ for both FKBP-DMSO and FKBP-DSS systems. The simulation box ultimately contained approximately 28K atoms, including all components of FKBP protein, DMSO, waters, and ions. To account for long-range electrostatics interactions, we calculated using the particle-mesh Ewald (PME) with a cutoff distance of 1.2 nm for both system. Similarly, the box contained approximately 27K atoms, including all components of FKBP protein, DSS, waters, and ions. Once the system was prepared for simulations, the remainder of the steps followed the detailed protocols outlined in Section II D of the main text.

#### T4 Lysozyme L99A-Benzene

MD simulation was initiated using the crystal structures of benzene bound to the T4 Lysozyme L99A (PDB ID: 1L83). Initially, before setting up the simulation box, we removed all existing water molecules

*

from the crystal structure. Subsequent setup steps were prepared using the CHARMM-GUI [1] solution builder module. This includes the addition of sodium ($Na^+$) and chloride ($Cl^-$) ions to achieve an ionic concentration of 100 mM, along with the addition of TIP3P water molecules [2] for solvation. During the solvation, a total of 27 $Na^+$ ions, 35 $Cl^-$ ions and 13k TIP3P water molecules were included in the simulation box. By solvating the structures in a TIP3P water box, one edge distance was extended to $\sim 77\AA$. The simulation box ultimately contained approximately 46K atoms, including all components of T4 Lysozyme L99A, benzene, waters, and ions. For long-range electrostatic interactions, we used the PME with a cutoff distance of 1.2 nm. The remainder of the simulation preparation followed the detailed protocols outlined in Section II D of the main text.

#### Abl kinase domain- imatinib

MD simulation was initiated using the crystal structure of imatinib bound to Abl kinase domain (PDB ID: 1OPJ). Mutant structures N368S and L364I were generated using the side-chain mutation module from MOE software. [5] Subsequently, the system was solvated using CHARMM-GUI [1] solution builder module. This involved adding sodium ($Na^+$) and chloride ($Cl^-$) ions to achieve an ionic concentration of 100 mM, and adding TIP3P water molecules [2]. During the solvation, By solvating the structures in a TIP3P water box, one edge distance was extended to $\sim 85\AA$. For long-range electrostatic interactions, the PME was used with a cutoff distance of 1.2 nm. The simulation box ultimately contained approximately 60 K atoms, including all components of Abl kinase protein, Imatinib, waters, and ions. The remainder of the simulation preparation followed the detailed protocols outlined in Section II D of the main text. These steps are followed from Ref. [4] supplementary information (SI).

### 2 SPIB accelerated time

In our SPIB-iMetaD protocol, we observed the reduction of the accelerated time at the completion of each round of simulation. This acceleration time refers to the Eq.1 shown in the main text. For direct comparison of dissociation rates, the reduction in acceleration time corresponding to each simulation round is shown in Fig. S1 and Fig. S2.

Figure S1 presents the SPIB-metadynamics iterations across three systems: (a) The FKBP-DMSO system underwent a total of 3 rounds of SPIB-metadynamics, achieving minimal accelerated time in round 2. Thereby, infrequent metadynamics (iMetaD) was performed upon completion of SPIB round 2. (b) The FKBP-DSS system completed a total of 2 rounds of SPIB-metadynamics, with the lowest accelerated time occurring in round 1. Thus, subsequent iMetaD was performed at SPIB round 1. (c) The T4 Lysozyme L99A-benzene underwent a total of 4 rounds of SPIB-metadynamics. Although the accelerated time slightly decreased in round 4 compared to round 3 of SPIB-metadynamics, the larger error bars at round 4 led to the selection of round three as the most preferred choice of RC for subsequent iMetaD simulations.

Figure S2 illustrates the SPIB-metadynamics iterations for Abl kinase domain with wild-type (WT) and two mutations bound with Imatinib: (a) The Abl kinase domain WT-Imatinib system underwent a total of 4 rounds of SPIB-metadynamics, reaching minimal accelerated time in round 3. Thereby, iMetaD was performed upon completion of SPIB round 3. (b) The Abl kinase domain with N368S mutation - Imatinib system completed a total of 3 rounds of SPIB-metadynamics, achieving the lowest accelerated time occurring in round 2. Thus, subsequent iMetaD was performed at SPIB round 2. (c) The Abl kinase domain with L364I mutation underwent a total of 3 rounds of SPIB-metadynamics. Despite a minor reduction in accelerated time in round 4 compared to round 3 of SPIB-metadynamics, the larger error bars at round 4 results made round 3 the preferred choice of RC for further iMetaD simulations.

### 3 p-value analysis

The Kolmogorov-Smirnov (KS) test is a crucial tool for evaluating whether sample data significantly differ from a theoretical distribution, particularly the exponential CDF. This test calculates the maximum distance between the empirical distribution of the sample and the exponential CDF. In particular, a low p-value ($< 0.05$) indicates significant differences between the empirical and exponential distributions such that we reject

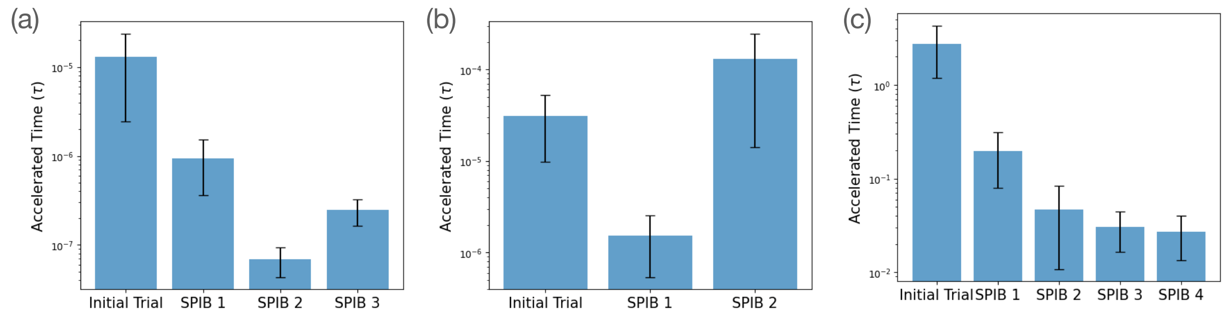


**Figure S1: SPIB accelerated time evaluation. (a) FKBP-DMSO system. (b) FKBP-DSS system. (c) T4 Lysozyme L99A-benzene system. Error bars denote the standard deviation from the mean value for each round of SPIB-metadynamics trajectories.**

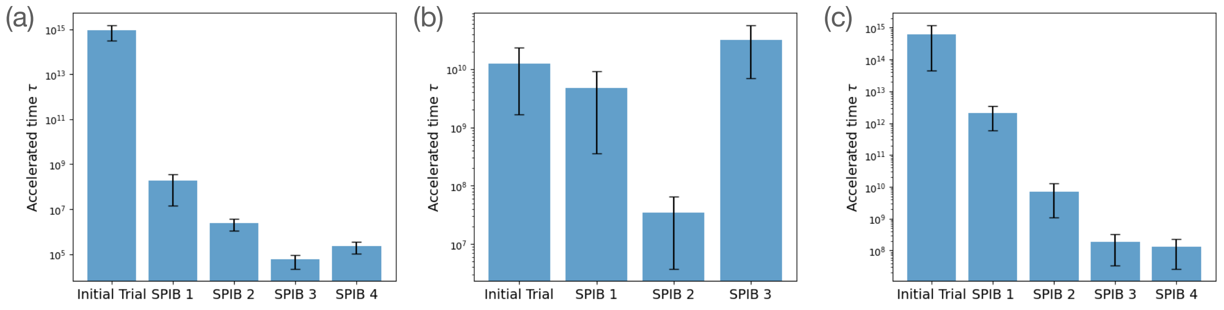


**Figure S2: SPIB accelerated time evaluations for Abl-kinase. (a) Abl-kinase WT with imatinib. (b) Abl-kinase mutant N368S with imatinib. (c) Abl-kinase mutant L364I with imatinib. Error bars denote the standard deviation from the mean value for each round of SPIB-metadynamics trajectories.**

the null hypothesis. This analysis, including curve-fit results and p-values, is presented in the following section.

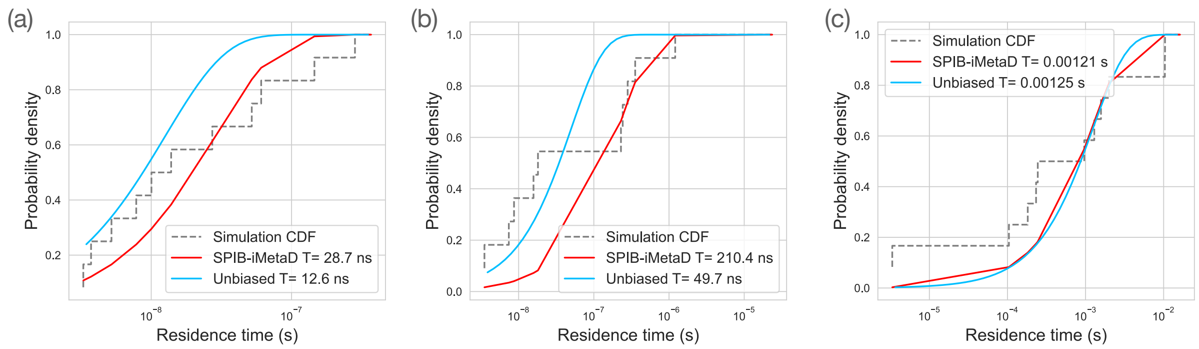


**Figure S3: FKBP-ligands and T4 Lysozyme L99A-benzene KS test and fitting results. (a) FKBP-DMSO ligand system curve fit results. The p-value was evaluated as 0.36. (b) FKBP-DSS ligand system curve fit results. The p-value was evaluated as 0.07. (c) T4 Lysozyme L99A-benzene system curve fit results. The p-value was evaluated as 0.38.**

### FKBP-ligands and T4 Lysozyme L99A-benzene

In the final stage of the protocol, 12 independent rounds of iMetaD were simulated for both the FKBP-DMSO and FKBP-DSS. The results of the curve fitting analysis and the KS test for these simulations are shown in Fig. S3 (a) and (b). Similarly, for the T4 Lysozyme L99A-benzene, the analysis involved conducting 10 independent rounds of iMetaD during the final stage. Subsequently, curve fitting was performed, and the results of the KS test are shown in Fig. S3 (c).

### Abl Kinase Domain- Imatinib dissociation pathway clustering and KS-test

The pathway clustering technique for analyzing dissociation pathways has been applied for the Abl kinase domain - imatinib system. As described in Ref. [4], the association of each pathway with varying dissociation time is important, potentially explaining the observed kinetic resistance in patients treated with Imatinib. Thus, we applied the pathway clustering approach exclusively for this system. While a more advanced approach has been proposed in other paper Ref. [3], we used simple metrics for pathway clustering. Specifically, we took the last 5 ns of the trajectory frame for the dissociation path analysis and distance calculations. We analyze two different paths, namely the C$\alpha$-helix and the front door pathways, the latter comprising both the P-loop and the hinge region. To determine the dissociation pathways of individual trajectories, we quantified three distinct distances: between the COM of Imatinib and the COMs of the hinge, P-loop, and C$\alpha$-helix regions. Following these measurements, we choose the shortest of the three distances as the dissociation pathway for each trajectory. The process of distance measurement and shortest distance selection was consistently applied across all trajectories. Subsequently, the trajectories were clustered based on these minimum distances using these defined proximity criteria.

Based on this clustering approach, we identified two primary dissociation pathways for all Abl-kinase on its WT and the mutants, depicted in Fig.S4. Particularly, Fig. S4 (a) shows the representative 'front door' pathways where imatinib dissociates near the P-loop and A-loop, highlighted in blue and magenta, respectively. Fig. S4 (b) shows the representative 'C$\alpha$-helix' pathways where imatinib dissociates closer to C$\alpha$-helix colored in green. This visual representation is used to help our understanding of the clustering mechanism. The summary of all these clustering results is presented in Table. S1.

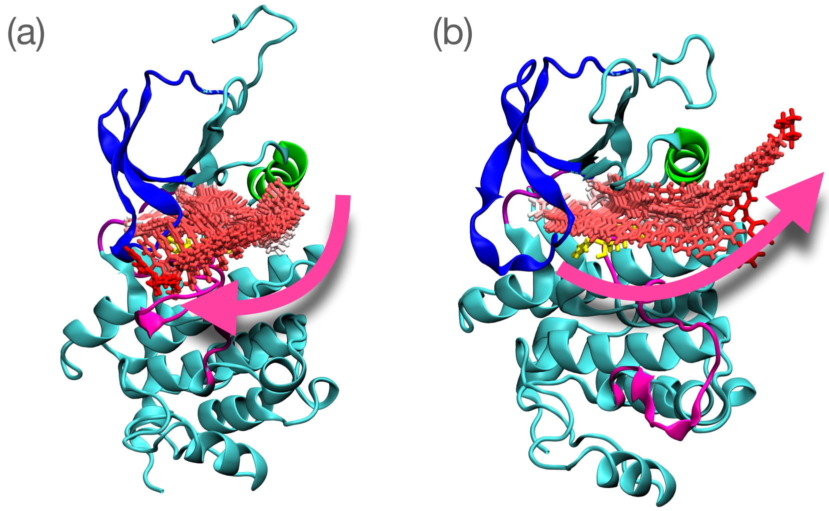


**Figure S4: Abl-kinase imatinib different dissociation pathways (a) Front door pathway: Imatinib dissociates closer to the P-loop (blue) and A-loop (magenta). (b) C$\alpha$-helix pathway: Imatinib dissociates closer to the C$\alpha$-helix (green) region.**

Based on the clustering analysis, we selected pathways that passed the p-value test. We did not choose the results with p-values less than 0.05, which was deemed less convincing. Consequently, we chose the front door to be the main pathway for both the wild-type and N368S mutant kinase and the C$\alpha$-helix pathway

| Abl Kinase | Dissociation path | Number of trajectories | Experimental time [s] | SPIB-iMetaD residence time [s] | P-value |
|---|---|---|---|---|---|
| WT | Cα-helix | 13/26 | 1200 | 40 | 0.03 |
| | Front | 13/26 | | 1150 | 0.07 |
| N368S | Cα-helix | 9/20 | 400 | 480 | 0.04 |
| | Front | 11/20 | | 330 | 0.19 |
| L364I | Cα-helix | 8/25 | 700 | 10000 | 0.06 |
| | Front | 17/25 | | 10700 | 0.01 |

**Table S1: Summary table for all Abl-kinase-imatinib experimental results (expected residence time) and simulation results (SPIB-iMetaD residence time) from our own semi-automated protocol approach depending on each dissociation pathway, including p-values.**

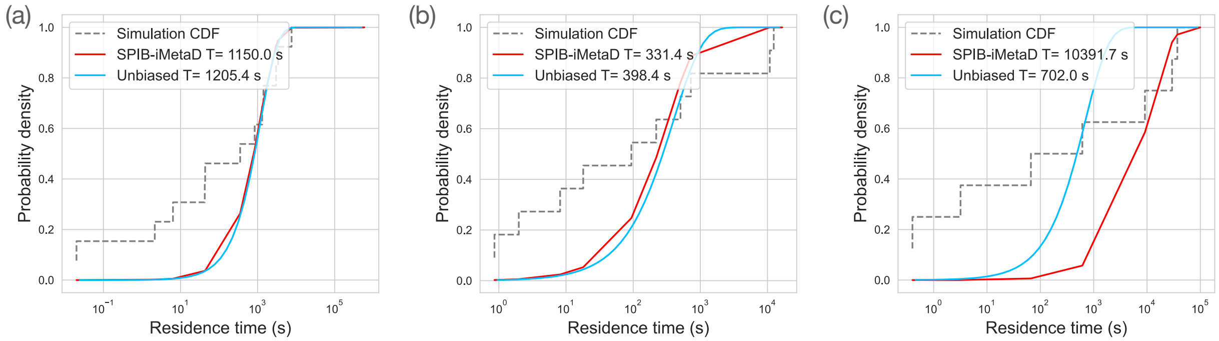


**Figure S5: Abl-kinase with Imatinib: (a) Abl-kinase wild-type (WT) system curve fit result. The p-value was evaluated as 0.07. (b) Abl-kinase mutant N368S system curve fit result. The p-value was evaluated as 0.19. (b) Abl-kinase mutant L364I system curve fit result. The p-value was evaluated as 0.06.**

for the L364I mutant. These results can be found in the main text, and curve fit and p-value results are presented in Fig. S5.
